# Protein interaction map of APOBEC3 enzyme family reveals deamination-independent role in cellular function

**DOI:** 10.1101/2024.02.06.579137

**Authors:** Gwendolyn M. Jang, Arun Kumar Annan Sudarsan, Arzhang Shayeganmehr, Erika Prando Munhoz, Reanna Lao, Amit Gaba, Milaid Granadillo Rodríguez, Robin P. Love, Benjamin J. Polacco, Yuan Zhou, Nevan J. Krogan, Robyn M. Kaake, Linda Chelico

**Affiliations:** Department of Cellular and Molecular Pharmacology, University of California San Francisco, San Francisco, CA, 94158, USA; Quantitative Biosciences Institute (QBI), University of California San Francisco, San Francisco, CA, 94158, USA; J. David Gladstone Institutes, San Francisco, CA 94158, USA; University of Saskatchewan, College of Medicine, Biochemistry, Microbiology & Immunology, Saskatoon, Saskatchewan, Canada; Centre for Commercialization of Regenerative Medicine (CCRM), 661 University Ave #1002, Toronto, ON M5G 1M1; Department of Medicine, Cumming School of Medicine, University of Calgary, 3330 Hospital Drive NW Calgary, AB T2N 4N1; Faculty of Medicine & Dentistry, Department of Medicine, TB Program Evaluation & Research Unit, University of Alberta, 11402 University Avenue NW, Edmonton, AB, T6G 2J3

## Abstract

Human APOBEC3 enzymes are a family of single-stranded (ss)DNA and RNA cytidine deaminases that act as part of the intrinsic immunity against viruses and retroelements. These enzymes deaminate cytosine to form uracil which can functionally inactivate or cause degradation of viral or retroelement genomes. In addition, APOBEC3s have deamination independent antiviral activity through protein and nucleic acid interactions. If expression levels are misregulated, some APOBEC3 enzymes can access the human genome leading to deamination and mutagenesis, contributing to cancer initiation and evolution. While APOBEC3 enzymes are known to interact with large ribonucleoprotein complexes, the function and RNA dependence is not entirely understood. To further understand their cellular roles, we determined by affinity purification mass spectrometry (AP-MS) the protein interaction network for the human APOBEC3 enzymes and map a diverse set of protein-protein and protein-RNA mediated interactions. Our analysis identified novel RNA-mediated interactions between APOBEC3C, APOBEC3H Haplotype I and II, and APOBEC3G with spliceosome proteins, and APOBEC3G and APOBEC3H Haplotype I with proteins involved in tRNA methylation and ncRNA export from the nucleus. In addition, we identified RNA-independent protein-protein interactions with APOBEC3B, APOBEC3D, and APOBEC3F and the prefoldin family of protein folding chaperones. Interaction between prefoldin 5 (PFD5) and APOBEC3B disrupted the ability of PFD5 to induce degradation of the oncogene cMyc, implicating the APOBEC3B protein interaction network in cancer. Altogether, the results uncover novel functions and interactions of the APOBEC3 family and suggest they may have fundamental roles in cellular RNA biology, their protein-protein interactions are not redundant, and there are protein-protein interactions with tumor suppressors, suggesting a role in cancer biology.

## INTRODUCTION

The APOBEC (apolipoprotein B mRNA editing enzyme, catalytic polypeptide-like) family of enzymes are single-stranded (ss) polynucleotide cytosine deaminases (1). In humans, there are 11 total members that are named after the first enzyme discovered, APOBEC1, which edits apolipoprotein B mRNA in addition to other mRNAs and ssDNA (2, 3). APOBECs that are primarily ssDNA deaminases have diverse roles in affinity maturation of antibodies (AID, activation induced cytidine deaminase), suppression of viral replication (APOBEC3 (A3) subfamily, A3A-H, excluding E), suppression of retrotransposons (APOBEC1, AID, A3s), and other yet to be characterized functions (APOBEC2, APOBEC4) (1). Many of the family members also have deaminase independent functions based on their ability to bind RNA and ssDNA with high affinity, such as blocking virus or retrotransposon polymerase progression (4–6). Through a hydrolytic reaction, APOBEC enzymes deaminate cytosine to form uracil (7) on transiently available ssDNA substrates, such as newly synthesized DNA, DNA being transcribed, or during DNA repair (8, 9). The cytosines deaminated are in a specific motif, such as 5’TTC (A3F) or 5’ATC (A3B) (10, 11). The fate of the resulting uracil is varied and may include acting as a template during downstream synthesis, which ultimately creates C to T transition mutations, or could result in the removal of the uracil by uracil DNA glycosylase, leaving an abasic site (9). This abasic site can be repaired with high or low fidelity, which may remove any effect of the uracil or cause the conversion of the C to A, G, or T (9). While the versatility of uracil in nucleating diverse downstream events is usually used as an advantage, some nuclear localized APOBECs can aberrantly deaminate genomic DNA during replication or transcription and this has been linked to ongoing mutagenic processes in tumors and cancer evolution (12). Some A3 enzymes can also promote genomic instability in the absence of deamination, but the mechanism is not known (13). There are multiple levels of control that attempt to suppress these “off-target” deaminations, such as cytoplasmic localization or binding to cellular RNA (14). Binding to cellular RNA results in APOBEC enzymes forming high molecular mass ribonucleoprotein (RNP) complexes that are localized to cytoplasmic RNA processing bodies.

Since their discovery over 20 years ago there have been essential functions described for many but not all of the APOBEC enzymes. Humans require APOBEC1 for proper lipid absorption by editing of the apolipoprotein B mRNA and deletions of APOBEC1 in mice cause lethal toxicity (15). AID is required for affinity maturation and class switching of antibodies and people born with genetic defects in AID are severely immunocompromised (1, 16). Although the A3 enzymes have essential roles in suppressing viral replication of retrotransposons and various viruses (e.g., Human immunodeficiency virus (HIV), Hepatitis B virus, Epstein Barr virus), many of them are thought to be functionally redundant (6, 17, 18). Thus, it is unclear why primates would maintain seven pro-mutagenic enzymes with four of the seven readily able to enter the nucleus (19). Three nuclear A3s, A3A, A3B, and A3H Haplotype I (Hap I), have been linked to mutagenesis and genomic instability in multiple cancer types (9, 13). At the population level, there is evidence of inactivation or reduced activity for two of the enzymes linked to cancer in humans. For example, there is a 90% chance of being homozygous for an *A3B* deletion in Oceanic populations (20). For other ethnicities it is 30% (20). In addition, A3H is highly polymorphic with seven main Haplotypes, with A3H Hap II, Hap V, and Hap VII being stable and active in cells (21, 22). However, the majority of humans encode either a hypo-active A3H (Hap I) that is ubiquitinated and degraded in cells, or an A3H (Hap III or Hap IV) that is ubiquitinated and degraded too quickly to observe catalytic activity (23). It has been hypothesized that these inactivating measures serve a protective role for the host genome.

We hypothesized that given the risk to the host for housing so many mutagenic enzymes, A3s must have additional functions that benefit humans that have not yet been discovered. In addition, it is likely that there are detrimental functions yet to be discovered that have supplied the evolutionary pressure for less activity of some A3 family members. Many proteins exist not alone, but within a protein interaction network to carry out their functions. Thus, we mapped the protein interaction network of eight A3 enzymes in order to identify connections to novel cellular pathways, functions and complexes. Using affinity purification mass spectrometry (AP-MS), we identified high confidence protein-protein interactions (PPIs) for A3A, A3B, A3C, A3D, A3F, A3G, A3H Hap I (A3H-I, hypo-active), and A3H Hap II (A3H-II, active) in the presence and absence of RNAse A to determine RNA-mediated and RNA-independent interactions. For both RNAse A treated and untreated conditions, we capture a number of A3-specific interactions, as well as a number of interactions that are shared across the A3 family. Among these are the Prefoldin (PFD) complex proteins (1–6) which specifically co-purify with A3B, A3D, and A3F under both RNAse A treated and untreated conditions. We demonstrate that PFD5 interaction with A3B reduces the PFD5 functional interaction with cMyc and stabilizes cMyc protein expression levels. Overall, we present the most comprehensive PPI network of the A3 family of enzymes to date. Functional enrichment analysis highlights novel cellular pathways and molecular functions that are likely deamination-independent, indicating that we have underestimated the physiological roles of A3 enzymes.

## EXPERIMENTAL PROCEDURES

### Experimental Design and Statistical Rationale

For each FLAG-tagged A3 bait (*e.g.* A3A, A3B, A3C, A3D, A3F, A3G, A3H-I, and A3H-II, in the not treated (NT) condition we performed 3 biological replicates. For the RNAse A treated conditions (+RNAse) the affinity purification experiments were performed as 3 (A3B, A3C, A3D, A3F and A3H-II) or 4 (A3A, A3G, A3H-I) biological replicates. No technical or process duplicates were performed for any sample collected. In total, we collected 51 experimental samples, 12 empty vector (EV) control samples (6 for NT, and 6 for +RNAse), and 11 FLAG-tagged green fluorescent protein (GFP) control samples (5 for NT, and 6 for +RNAse) (**Table S1**). The number of EV and GFP controls were selected based on the total number of independent AP-MS sample preparations, such that samples purified on separate days were represented by an EV and GFP control. EV controls were used for scoring by SAINTexpress (v 3.6.3) (24). GFP controls were scored by SAINTexpress as baits and used to verify purification background proteins were effectively filtered out of the high confidence PPI data. All A3 baits, EV, and GFP controls were analyzed by CompPASS (25) for increased confidence in identifying PPIs from purification background. Immunoblotting for co-immunoprecipitation (co-IP) experiments and quantification were performed in triplicate.

### Expression Constructs

All A3 expression constructs were obtained from the NIH HIV Reagent program except A3A, A3H-I, and A3H-II. The A3A and A3H-I cDNA were purchased and cloned into pcDNA with a 3×HA tag. A3H-II was created using site directed mutagenesis (26). The A3B plasmid obtained from the NIH HIV Reagent program (ARP-11090) was found to contain mutations which were corrected by site directed mutagenesis to match NCBI Accession AY743217. The NIH HIV Reagent program catalog numbers for the other plasmids were: A3C (ARP-10101, subcloned into pcDNA with a 3×HA tag), A3D (ARP-11433, subcloned into pcDNA with a 3×HA tag), A3F (ARP-10100, subcloned into pcDNA with a 3×HA tag), A3G (ARP-9952). Reading frames from these pcDNA backbones with 3×HA tags were amplified and inserted into pcDNA4/TO (Invitrogen) containing a C-terminal 3×FLAG affinity tag using Gibson assembly. All sequences were verified and matched against A3 reference sequences to ensure correct haplotype sequences were used and no mutations were introduced. We later discovered a point mutation in the A3C construct after initial AP-MS experiments were performed. It was unclear when this mutation was introduced, therefore the resulting samples were removed from the dataset, the mutation was corrected by site directed mutagenesis, and the corrected A3C plasmid was used for additional experiments.

### Cell Culture, transfection, and cell harvest for immunoprecipitation

HEK293T cells were obtained from the UCSF Cell Culture Facility (https://cgec.ucsf.edu/cell-culture-and-banking-services), and cultured in Dulbecco’s Modified Eagle’s Medium (4.5 g/L glucose, 0.584 g/L L-glutamine, and 3.7 g/L NaHCO_3_, DMEM) supplemented with 10% Fetal Bovine Serum (Gibco, Life Technologies), 1% Penicillin-Streptomycin (Corning), and 1% Sodium Pyruvate. Cells were grown and maintained at 37°C in a humidified atmosphere of 5% CO_2_ in T175 flasks (Corning). For each immunoprecipitation, HEK293T cells were plated in 2×15-cm dishes at 1×10^7^ cells per plate. After 20-24 hrs of recovery, the HEK293T cells were transfected with up to 15 μg of the individual 3×FLAG A3 construct or GFP control DNA. All DNA mixtures were complexed with PolyJet In Vitro DNA Transfection Reagent (SignaGen Laboratories) at a 1:3 μg:μl ratio (plasmid:transfection reagent) according to the manufacturer’s recommendations. Plasmids and PolyJet mixtures were each separately diluted in 0.5mL DMEM, combined and vortexed before incubating 20 minutes at room temperature before adding dropwise to cells. Transfected cells were grown for 40-48 hrs, then the media was removed and the cells were dissociated from the 15-cm dishes using room temperature Dulbecco’s Phosphate Buffered Saline without calcium and magnesium (D-PBS) supplemented with 10 mM EDTA. Cells from each pair of transfected dishes were combined and pelleted by centrifugation at 200xg for 5min at 4°C and washed with 10 ml D-PBS. Cells were pelleted by centrifugation, divided into two equal aliquots, and recollected by centrifugation at 500xg for 5 min at 4°C. Each of the baits and controls was individually prepared with at least three biological replicates.

### Cell lysis and anti-FLAG immunoprecipitation

Cell pellets were lysed on ice in 1mL of freshly prepared cold non-denaturing lysis buffers without or with or RNAse A added (IP buffer [50mM Tris-HCl, pH 7.4 at 4°C, 150mM NaCl, 1mM EDTA], supplemented with 0.5% Nonidet P 40 Substitute (NP40; Fluka Analytical), and cOmplete mini EDTA-free protease and PhosSTOP phosphatase inhibitor cocktails (Roche)). RNAse A lysis buffer was prepared from IP buffer but is supplemented with 80 μg/mL of RNase A (Qiagen). Cells were gently resuspended and incubated for 30 minutes rocking on a tube rotator at 4°C. Lysates were clarified by centrifugation at 3,500xg at 4°C for 20 minutes, supernatants were collected in fresh 1.5mL protein lo-bind tubes (Axygen) and cell debris discarded. A small amount of each lysate (50 μL) was reserved to monitor bait protein expression and cell lysis by immunoblotting and silver staining. Anti-FLAG M2 magnetic beads (40 μL slurry; Sigma-Aldrich) were initially washed twice in 1.0 mL IP buffer supplemented with 0.05% NP40 and then kept in 0.3 mL IP buffer supplemented with 2 μg 1xFLAG peptide. Cell lysates were combined with washed anti-FLAG M2 magnetic beads and incubated at 4°C for 2 hrs on a tube rotator. After binding, the flow thru was collected and the beads were washed three times with 1 mL wash buffer (IP buffer with 0.05% NP40). Beads were transferred to a clean tube for a final wash in 1 mL of IP buffer. Bound proteins were eluted by gently agitating the FLAG beads with 30 μL 0.05% RapiGest SF Surfactant (Waters Corporation) in IP buffer at room temperature for 30 min using a vortex mixer. Lysates and eluates were resolved on 4-20% Criterion protein gels (Bio-Rad Laboratories) to assess FLAG-tagged protein expression and immunoprecipitation by Immunoblot and silver stain (ThermoFisher Scientific) respectively. For each immunoprecipitation, 10 μL of eluate was submitted for protein digestion and analysis by liquid chromatography tandem mass spectrometry (LC-MS/MS).

### Reciprocal immunoprecipitation of endogenous PUS7 protein under NT and +RNase conditions

Endogenous immunoprecipitation of PUS7 was performed using the KingFisher Flex (KFF) Purification System (Thermo Fisher Scientific). Beads and buffers (indicated below) were dispensed into KingFisher 96-well deep-well plates or microplates as appropriate and placed on ice until loaded onto the instrument for automated processing as follows: Antibodies (α-PUS7 or IgG1 isotype control were added to (plate 1) 0.5 mL cell lysate and brought up to 0.75 mL with Lysis Buffer. After incubating on KFF for 2 hrs, pre-equilibrated Pierce Protein A/G magnetic beads (originally 12.5 μL slurry) were added to lysate-antibody and incubated for an additional 2 hrs. Protein-bound beads were washed four times (plates 2-5) with 1.0 mL IP Buffer supplemented with 0.05% NP40 and eluted in (plate 6) 50 μL 0.05% RapiGest in IP Buffer. The KFF is operated in a cold room to maintain a 4°C temperature during immunoprecipitation; however, elutions were performed with the heat block pre-heated to 23°C. Automated protocol steps were performed using the slow mix speed and the following mix times: 2 hrs for binding steps, 2 m for wash steps, and 35 min for the elution step. Two rounds of bead collection (five 30 sec bead collection times) were used at the end of each step before transferring beads to the next plate. After the elution step, the instrument was paused for 2 min, to allow beads to settle prior to bead collection.

### SDS-polyacrylamide gel electrophoresis (SDS-PAGE), silver stain, and immunoblot analysis of affinity purified proteins

Proteins were separated by 4-20% SDS-PAGE and either silver stained (eluate only; ThermoFisher Scientific) according to manufacturer’s protocols, or transferred to PVDF membranes at 0.25 A for 1.5 hrs at 0°C. Transferred PVDF membranes were blocked in 5% milk powder in 0.2% Tween-TBS overnight at 4°C and immunoblotted with mouse anti-FLAG-HRP (Sigma) conjugated primary antibody, rabbit α-ELAC2 (RNZ2) polyclonal antibody (Proteintech, 10071-1-AP, 1:2,000) and mouse α-PUS7 monoclonal antibody (OriGene, OTI4C6; 1:1,000) followed by goat α-rabbit, and α-mouse secondary antibodies (BioRad; 1:10,000), respectively. Pierce ECL Western Blotting Substrate (Thermo Scientific) was used to detect bands with Hyperfilm ECL film (Amersham).

### Protein digestion and peptide desalting

RapiGest MS-safe eluted proteins (10 μL) were reduced in 2 M urea, 10 mM NH_4_HCO_3_, and 2 mM DTT at 60°C for 30 minutes with constant shaking. Samples were then alkylated in the dark at room temperature with 2 mM iodoacetamide for 45 minutes. Reduced and alkylated proteins were digested with 80 ng trypsin overnight at 37°C. Peptides were acidified with formic acid (pH < 3) and then desalted using C18 ZipTips (Millipore) according to the manufacturer’s protocols. Desalted peptides were centrifuged in a speedvac to dry and stored at −80°C.

### Mass spectrometry data acquisition

Dried peptide samples were resuspended in 2% acetonitrile, 0.1% formic acid solution and analyzed by LC-MS/MS using an Easy-nLC 1000 (Thermo Fisher Scientific) coupled online to an Orbitrap Elite Hybrid Mass Spectrometer with ETD (Thermo Fisher Scientific). Peptides were separated on a 75 μm × 25 cm fused silica IntregraFrit capillary column (New Objective) packed in-house with 1.9-μm Reprosil-Pur C18 AQ reverse-phase resin (Dr. Maisch-GmbH). Peptides were eluted at a 300 nL/min flow rate over a 60 minute gradient: 5% B for 1 min, 5%–30% B in 50 min, 30%–95% B in 5 minutes, and 95% B for 4 min (mobile phase buffer A: 100% H_2_O/0.1% FA; mobile phase buffer B: 100% ACN/0.1% FA). Each immunoprecipitation sample was run once with a clean gradient run between each individual bait samples, and all samples were randomized in the queue to reduce carry-over effects. Data was collected on the Orbitrap Elite in positive ion mode with MS1 detection in profile mode in the orbitrap (150–1500 m/z, 120,0000 resolution, AGC target of 1×10^6^, max injection time of 100 ms). MS2 fragmentation was performed on charge states 2+ and above with normalized collision energy set at 35% with a 20 second dynamic exclusion after a single selection (tolerance of 10 ppm). MS2 data was collected in the ion trap (ion count target 10^4^, max injection time of 50 ms). For full description of all LC, MS acquisition, and tune parameters see **Table S2**.

### Peptide and protein identification and high confidence PPI scoring

All raw data files were searched with MaxQuant (27) (v 1.6.3.3) against the human proteome (SwissProt canonical protein sequences-20393 entries, updated October 09, 2018) concatenated with a fully randomized decoy database, using a 0.01 peptide and protein false discovery rate. The following default MaxQuant parameters were used: 1) digestion mode was set to specific and Trypsin/P was selected with 2 max missed cleavages; 2) Carbamidomethyl (C) was selected as the only fixed modification; 3) Oxidation (M) and Acetyl (Protein N-term) were selected as the variable modifications with 5 max number of modifications; and 4) precursor and fragment mass tolerance were set to 20 ppm and 0.5 Da respectively. In addition label-free quantification was turned on, with match between runs set to 0.7 min. For each bait in each condition, PPIs were determined by scoring with SAINTexpress (v 3.6.3) (24) and CompPASS (25) with both the NT and +RNAse EV samples being combined to use as controls. Since we collected an EV and GFP control each time an individual replicate was generated, and not every replicate included all baits, more controls were collected than individual A3 baits. As only 3-4 individual samples collected per bait per condition, we randomly selected 12 total EV and 11 GFP controls for SAINT scoring, with EV acting as the SAINT control, and GFP acting as a bait for further confidence filtering. The metadata describing the files associated with each of the biological replicates for each bait in each condition can be found in **Table S1**. We applied a two step filtering strategy to determine the final list of reported high confidence interactors which relied on two different scoring stringency cutoffs. In the first step, for each bait and condition, an identified protein must have a SAINTexpress Bayesian False Discovery Rate (BFDR) < 0.05. In the second step, an identified protein was considered a high confidence interactor for that bait and condition, if it was in the 0.9 percentile of the CompPASS wd percentile per bait score and removing any preys from the list which are also called hits in the appear in GFP control (which was treated as a bait during scoring). High confidence interactions for each bait were mapped in separate condition specific networks, as well as a combined network, and visualized with cytoscape (28). All mass spectrometry data files (raw and MaxQuant search results), as well as associated metadata, and SAINTexpress scored data files, have been deposited to the ProteomeXchange Consortium (29) via the PRIDE (30) partner repository.

### Differential interaction score (DIS) and MSStats analysis

A differential interaction score (DIS) was computed for all high confidence interactions identified for any A3 bait in either NT or +RNase conditions (described in (31–33)). The DIS is calculated as the difference between the NT and +RNase interaction scores for a given bait with the interaction score being the average of 1 - the SAINTexpress BFDR and CompPASS wd percentile per bait. In this way, a DIS near 0 indicates an interaction that is confidently identified in both NT and +RNAse conditions, while a DIS near −1 or +1 indicates that interaction is specific to NT or +RNAse conditions respectively. MSstats analysis was used to quantitatively compare and measure significant differences in protein abundance between NT and +RNAse conditions using default parameters for MSstats and adjusted P values (Student’s t test and Benjamini–Hochberg correction) (34). This quantification was performed using artMS, acting as a wrapper for MSstats, utilizing the function artMS::artmsQuantification with default settings.

### A3 domain sequence alignment and PPI similarity analysis

For each A3 protein, the HMMER (v 3.3.1, http://hmmer.org/) (35) tool hmmmscan was used to match against Pfam (36) A domains in order to define double APOBEC domain boundaries. Specifically, matches to Pfam domains of the following terms were selected: “NAD2”, “APOBEC2”, “NAD1”, APOBEC_N”, and “APOBEC_C”. The two domain A3s were then split into two by dividing at the midpoint between the largest first and second domain matches found by hmmscan. An all-by-all pairwise identity between domains was calculated using BLAST (v 2.11.0+) (37) with Smith-Waterman traceback enabled. Multiple sequence alignment and clustered trees were calculated by clustal-w (v 2.1) (38). Pairwise comparisons of the high confidence PPIs were calculated and visualized using the online ProHits-viz analysis and visualization tools. Bait comparisons were performed for each condition separately, as well as a full set, with the abundance set to the average of the spectral count (AveSpec). Session files for each comparison (NT_244_bait_session.json; RNAse_244_bait_session.json; ALL_244_bait_session.json) are provided in the supplement and can be uploaded to the Prohits-viz visualization tool (39) to display matrices and analysis settings.

### Functional over-representation/enrichment analysis

The high confidence PPIs for each bait under each condition were combined to test for A3 specific functional enrichments of Gene Ontology (GO), KEGG, and canonical pathway terms. Over-representation analysis was performed using Metascape Express Analysis (https://metascape.org) with default parameters. The top significant terms (based on cumulative hypergeometric p-values and enrichment factors) were identified, refined to non-redundant terms, and clustered hierarchically. In addition to the bait specific enrichments, PPIs for each condition were combined (all baits) to identify any major functional differences between both conditions. Enrichments are visualized by heatmaps that were automatically generated in the analysis report by Metascape. Subcellular localization of PPIs for A3B and A3F were determined by manually curating the subcellular localization information from https://www.uniprot.org/. The percent of prey localized to the nucleus was calculated for A3B and the percent of prey proteins localized to the cytoplasm was calculated for A3F.

### Transfections in HEK293T cells for reciprocal and experimental anti-FLAG immunoprecipitation

HEK293T cells were maintained in Dulbecco’s modified Eagle medium with Fetal Bovine Serum to a final concentration of 10%. Cells were grown and maintained at 37°C in a humidified atmosphere of 5% CO_2_. HEK293T cells were seeded in a 75 cm^2^ tissue culture flask at 2×10^6^ cells per flask and 16 to 24 hrs later, plasmids were transfected using GeneJuice (MilliporeSigma/ Novagen) transfection reagent as per the manufacturer’s protocol. The following C-terminally 3×HA tagged constructs in pcDNA were used as indicated in the figures: A3B, A3D, A3F, A3G; C-terminally FLAG tagged constructs in pcDNA: PFD3 (GenScript, NM_003372), PFD5 (GenScript, NM_002624); and cMyc (GenScript, NM_002467) in pcDNA. After 24 hrs, where indicated 12.5 μM MG132 was added for 16 hrs. Then, 40 hrs after the transfection, cells were washed with 1x PBS and lysed using 1x Lysis Buffer (50mM Tris, 1% Nonident-P40, 0.1% sodium deoxycholate, 150mM NaCl, 10% glycerol, 50 mM sodium fluoride, and cOmplete mini EDTA-free protease inhibitor). Protein concentrations were estimated using Bradford assay. RNAse A at a final concentration of 50 μg/mL was used for the required experiments. Subsequently, anti-FLAG M2 magnetic beads (Sigma) were added to the lysates and incubated for 2 hrs at 4 °C on a rotator. Co-IP was performed as per the manufacturer’s protocol, but using Lysis Buffer for washes. Laemmli buffer without reducing agent was used to elute proteins. Samples were reduced with 5% 2-mercaptoethanol and then resolved by SDS-PAGE before transferring to a nitrocellulose membrane. The 3×HA tagged A3B, A3D, A3F, A3G were detected using anti-HA rabbit antibody (1:1,000-5,000, Sigma), FLAG tagged PFD3 and PFD5 were detected using anti-FLAG rabbit antibody (1:5,000, Sigma), FLAG tagged PFD5 in combination with A3B and cMyc was detected using anti-FLAG mouse antibody (1:1,000, Sigma) and anti-c-Myc rabbit antibody clone 7E18 (1:1000, Sigma). Secondary detection was performed using Licor IRDye antibodies IRDye 680-labeled anti-rabbit. Secondary antibodies were used at 1:10,000.

### Transfections in HEK293T cells for detection of A3B, PFD5, and cMyc

HEK293T cells were plated in a 6 well plate at 3×10^6^ cells per well and GeneJuice transfection reagent was used as per the manufacturer’s protocol. The following plasmids were transfected (100 ng each): pcDNA A3B-3×HA, pcDNA PFD5-FLAG, and pcDNA cMyc. Empty pcDNA was used to equalize transfection amounts to 300 ng where needed. Transfection media was replaced the day after transfection and MG132 was added to a final concentration of 12.5 μM for 16 hrs. Then, 40 hrs after the transfection cells were washed with 1x PBS and lysed in 2x Laemmli Buffer. A Lowry assay (Sigma - total protein kit) was used to determine protein concentrations before resolving samples on an SDS-PAGE gel and transferring to nitrocellulose membrane for immunoblotting. The A3B-3×HA was detected using anti-HA rabbit antibody (Sigma), FLAG tagged PFD5 was detected using anti-FLAG mouse antibody (Sigma) and cMyc using anti-c-Myc rabbit antibody clone 7E18 (Sigma). Loading control for cell lysate, α-tubulin, was detected using an anti-α-tubulin mouse antibody (Sigma). Secondary detection was performed using Licor IRDye antibodies IRDye 680-labeled anti-rabbit and IRDye 800-labeled anti-mouse. Primary antibodies were used at 1:5,000 and secondary antibodies were used at 1:10,000. Immunoblots were quantified using Image Studio software with normalization of each experimental lane to its respective anti-tubulin band.

### Transfection and immunoblotting for MCF7 cells

Six-well plates were seeded with MCF7 cells at 3×10^5^ cells per well and GeneJuice transfection reagent was used as per the manufacturer’s protocol. Cells were maintained in Eagle’s Minimum Essential Medium with 0.01 mg/mL human recombinant insulin and Fetal Bovine Serum to a final concentration of 10%. Cells were grown and maintained at 37°C in a humidified atmosphere of 5% CO_2_. C-terminally tagged A3B-3×HA construct in pcDNA was used for transfections (0-300 ng) using GeneJuice transfection reagent as per the manufacturer’s protocol. After 48 hrs, cells were washed with 1x PBS and lysed using 2x Laemmli buffer. Protein concentrations were estimated using a Lowry assay (Sigma – total protein kit). A3B was detected using an anti-HA rabbit antibody (Sigma), endogenous cMyc and PFD5 were detected using rabbit anti-c-Myc antibody clone 7E18 (Sigma) and mouse anti-prefoldin 5 antibody (Santa Cruz), respectively. Loading control for cell lysate, α-tubulin, was detected using an anti-α-tubulin mouse antibody (Sigma). Secondary detection was performed using Licor IRDye antibodies IRDye 680-labeled anti-rabbit and IRDye 800-labeled anti-mouse. Primary antibodies were used at 1:1,000 and secondary antibodies were used at 1:10,000. Immunoblots were quantified using Image Studio software with normalization of each experimental lane to its respective anti-tubulin band.

### siRNA knockdown of PFD5 in A3B-FLAG expressing MCF7 cells

The doxycycline (dox) inducible A3B-FLAG expressing stable MCF7 cells have been previously reported (40). Silencer^TM^ siRNAs were ordered against PFD5 and a scramble of the siRNA (ThermoFisher Scientific). The Silencer Select Transfection Protocol from the manufacturer was followed using Lipofectamine RNAiMAX as a transfection reagent. Six-well plates were seeded with MCF7 cells at 3×10^5^ cells per well and siRNA were transfected. Twenty-four hours after the transfection, dox was or was not added to the wells. Forty-eight hours after addition of dox, cells were washed with 1x PBS and lysed using 2× Laemmli buffer. Protein concentrations were estimated using a Lowry assay (Sigma – total protein kit). A3B was detected using an anti-FLAG mouse antibody (Sigma), endogenous cMyc and PFD5 were detected using rabbit anti-c-Myc antibody clone 7E18 (Sigma) and mouse anti-prefoldin 5 antibody (Santa Cruz) respectively. Loading control for cell lysate, α-tubulin, was detected using an anti-α-tubulin mouse antibody (Sigma). Secondary detection was performed using Licor IRDye antibodies IRDye 680-labeled anti-rabbit and IRDye 800-labeled anti-mouse. Primary antibodies were used at 1:500 (cMyc, FLAG, PFD5) or 1:1000 (α-tubulin) and secondary antibodies were used at 1:5,000. Immunoblots were quantified using Total Lab Quant v12.5 with normalization of each experimental lane to its respective anti-tubulin band.

## RESULTS

### Global analysis of A3 protein-protein interactions (PPIs)

Here we use a label-free FLAG-based AP-MS approach to characterize the RNA -dependent and -independent PPI network for human A3 proteins (**Figure 1A**). To this end, we transiently transfected HEK293T cells with C-terminally 3×FLAG tagged A3 proteins (*i.e.* A3A, A3B, A3C, A3D, A3F, A3G, and two haplotypes of A3H: A3H-I and A3H-II), or FLAG tagged negative controls (*i.e.* GFP-FLAG, empty vector (EV)) (**Figure S1**). A3A is the only A3 family member expressed in myeloid lineage cells and the others are primarily expressed in CD4+ T cells at different levels (41, 42). During viral infection and cancer expression of some A3s has been identified in other cell types, but there is no consistent cell type where their expression is found (18, 43–47). Given these reasons as well as the frequency of HEK293Ts being used as model cells for PPI network analyses, we used HEK293T cells to compare interactions across A3 family members. Negative controls and A3-FLAG proteins were affinity purified by anti-FLAG immunoprecipitation in biological triplicate from HEK293T cell lysates that were either treated with RNAse A (+RNAse) or not treated (NT) (**Figure S2**). Purified samples were digested with trypsin, and resulting peptides analyzed by liquid chromatography tandem mass spectrometry (LC-MS/MS). Data was searched using MaxQuant (27) (**Table S3-4**), and high confidence PPIs were differentiated from background using a two-step filtering approach with the SAINTexpress (24) and CompPASS (25) scoring algorithms.

**Figure 1.**
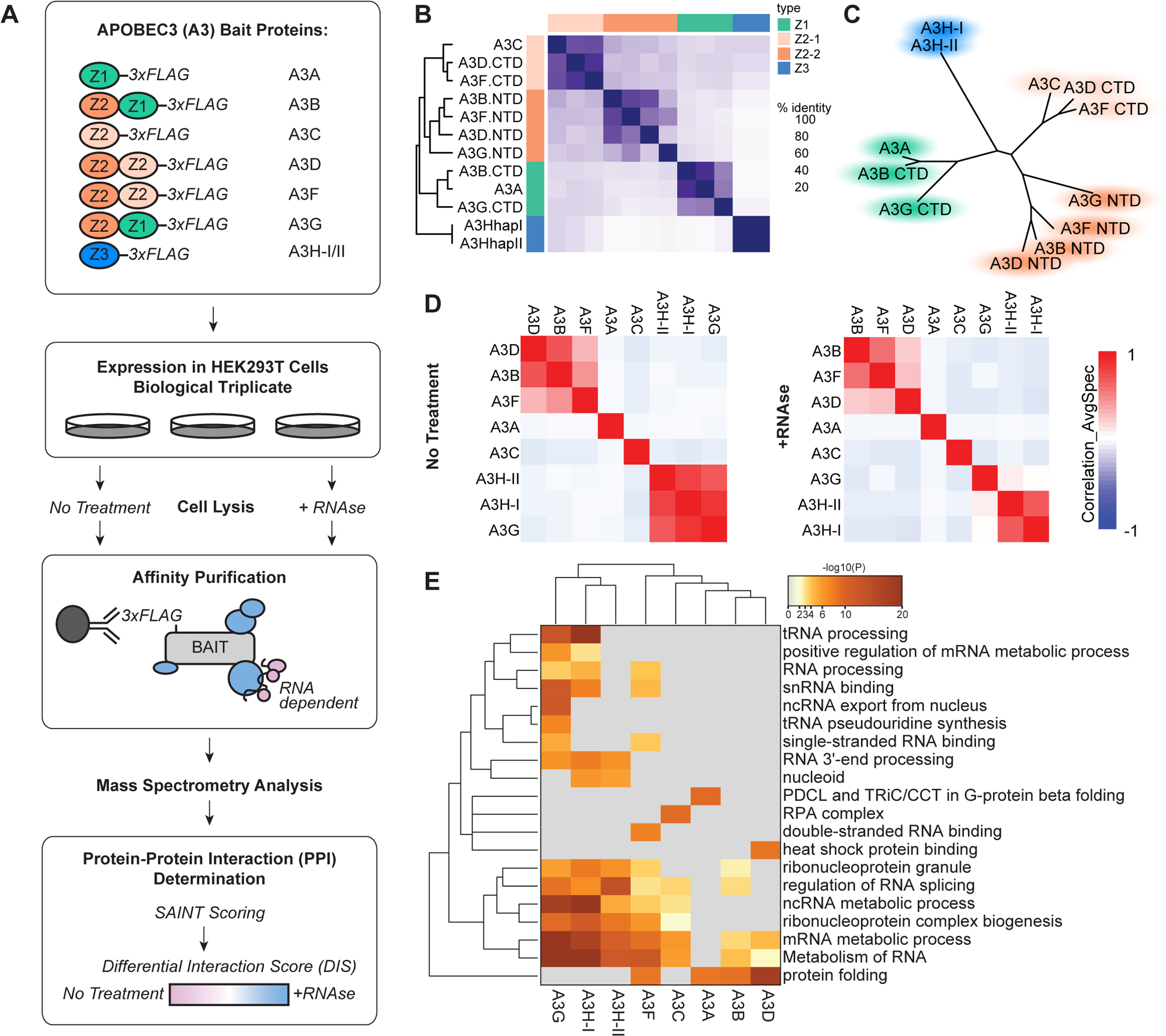
AP-MS analysis of the A3 family proteins identifies shared and specific RNA-dependent and -independent PPIs. **(A)** In the AP-MS workflow described in this study, HEK293T cells were transiently transfected with C-terminally 3×FLAG-tagged A3 proteins in biological triplicate and purified with or without RNAse A treatment. Purified proteins were trypsin digested and analyzed by LC-MS/MS. Peptides and proteins were identified by MaxQuant (27), high confidence PPIs were determined by SAINT and CompPASS, and RNA-dependent and independent PPIs were characterized using a Differential Interaction Score (DIS) calculation. The cartoon representation at the top shows each of the 3×FLAG tagged A3 proteins with the Z1 domain in green, the Z2 domain in orange, and Z3 domain in blue. **(B)** Sequence identity matrix demonstrating pairwise similarity in A3 domains. **(C)** Clustered tree diagram of the similarity of A3 protein domains. **(D)** Correlation plot of SAINT scored proteins identified in A3 pull-downs with no treatment (left) and RNAse A treatment (right); performed by Prohits-viz (39)**. (E)** Heatmap of the top functional enrichment analysis of high confidence PPIs per each bait. For the full list of enrichments see **Table S7**.

Using SAINTexpress we first filtered out any protein with a BFDR ≤0.05 (**Table S5, Figure S3**) and then removed any preys from the list which were also called hits in the GFP control (one prey), resulting in the identification of 744 interactions between the eight A3 bait and 292 prey proteins. This included 391 interactions in RNAse+ (blue edges) and 353 interactions in NT (red edges) conditions; of these 744 interactions, 326 bait-prey pairs (163 interacting proteins) were captured in both conditions. In total, 129 proteins in this network were identified as interacting with at least two A3 bait proteins in at least one condition, while 163 proteins were found to pull-down specifically with one A3 protein in one or both conditions. SAINTexpress accounts for reproducibility, abundance, and specificity against a control (in this case, the empty vector), regardless of the other baits and therefore avoids penalizing proteins that are identified across multiple baits or within different conditions, which is good for a network where sequence similarity between baits and shared nucleic acid binding function in cells could result in overlapping interactors. However, given the density and size of the SAINTexpress network, we used CompPASS wd scores (0.9 percentile per bait) as a second filter to focus on the highest confidence interactions, and clarify which hits are more likely to be strong A3 and condition-specific interactors; these are highlighted as darker cyan in **Figure S3** (**Table S5**). In total, we captured 143 and 153 high confidence PPIs between the eight A3 protein baits under NT and +RNAse conditions respectively, with 60 PPIs being identified in the A3-PPI network under both conditions (**Figure S4A**). In comparison to the SAINTexpress network, this reduced the total prey count by 132 and corresponded to 102 and 101 total high confidence interacting proteins identified in NT and +RNAse conditions respectively, with 48 being shared across conditions (**Figure S4B**). The proteins identified in each condition for each individual A3 bait can differ fairly significantly, thus demonstrating the importance of performing these analyses for each bait under both conditions (**Figure S4C**).

To see if protein domain structure and sequence similarity correlated with high confidence A3 co-purifying proteins in NT and +RNAse conditions, we calculated the pairwise correlation and clustered in a correlation matrix. The A3 proteins are a family of single (A3A, A3C, A3H) and double (A3B, A3D, A3F, A3G) Zinc coordinating domain (Z domain) proteins, with the protein domains all sharing a core structure of a five-stranded β-sheet surrounded by six α-helices (48). The double domain proteins were considered as two entities, the N-terminal domain (NTD) and the C-terminal domain (CTD). Sequence alignment of the amino acids in each domain and similarity analysis clusters these protein domains into four groups: Z1 domains (A3A, A3B-CTD, and A3G-CTD), the Z3 domain (A3H), and two groups of Z2 domains (1-A3C, A3D-CTD, A3F-CTD; 2-A3G-NTD, A3F-NTD, A3B-NTD, A3D-NTD) (**Figure 1B and 1C**) (49, 50). However, these amino acid similarities did not always correlate with the clustering of shared PPIs (**Figure 1D**). Further, even though some of the A3 proteins have similar documented functions, such as the anti-HIV activity of A3F, A3G, and A3H-II, most have diverse PPIs (**Figure 1D**). Comparing single domain A3A, A3C, and A3H proteins, we do not see high levels of overlapping interactions between them and other A3 proteins. A3A does not share high confidence interactors with the other A3 proteins, and its interactors are largely not sensitive to RNAse A treatment (**Figure S5**). Similarly to A3A, A3C does not have a high degree of shared high confidence interactions and only shares one interaction with any other A3 protein (**Figure S4C**). The two A3H haplotype baits, A3H-I and A3H-II, share a number of interactors and cluster together in both NT and +RNAse conditions. Notably, under NT conditions these A3H proteins also cluster with and share interactors with A3G, though this correlation is lost in the +RNAse condition. Given A3H is a single domain, whereas A3G is a double domain A3 protein, and the low sequence homology between the domains, the overlapping proteins likely reflect a functional similarity. Given these interactions are sensitive to RNAse A, it is probable that these interactions are mediated by RNA. The double domain proteins A3B, A3D, and A3F cluster together under both NT and +RNAse conditions, indicating shared interactions. Based on the domain structure and homology of these proteins (**Figure 1B**), the NTDs of A3B, A3D, and A3F cluster together and it is feasible that the shared interactions are engaging similar binding surfaces on each protein through their NTD. Notably, when comparing correlation across all baits and conditions, this trend remains (**Figure S5**).

All PPI data was collected by affinity purification in which cells are lysed in non-denaturing buffer prior to purification, thus disrupting cellular compartmentalization, which could allow otherwise location-dependent interactions to occur. Most A3 proteins are found throughout the cell (19) and all A3s except A3A have been previously associated with ribonucleoprotein complexes in the cytoplasm, which is consistent with our PPI dataset (14, 51–54). Additionally, although A3D and A3G were thought to be only cytoplasmic, several more recent reports have shown that they are also found in the nucleus (46, 55, 56). Only A3B has a nuclear localization signal that places the majority of A3B in a single compartment, but similar to other A3s, there are conditions where it becomes relocalized to the cytoplasm (53, 57). A3F has largely been identified in the cytoplasm except during HIV-1 infection (53). Since A3B and A3F are the most discretely localized bait proteins, we used their PPI datasets to test the specificity of the prey proteins to cellular localization. We found that for A3B PPIs, 62.5% were proteins that had subcellular localization reported in the nucleus (**Table S6**). Conversely, for A3F, we found that 77% of the PPIs were proteins that had subcellular localization reported in the cytoplasm, indicating that a number of compartment specific interactions were maintained in our PPI dataset (**Table S6**).

We performed functional enrichment analysis for each bait in order to further characterize the high confidence PPIs pulled down by each A3 bait. Here we were primarily interested in characterizing bait-specific and shared interactions, and since some shared interactors were conditional for some baits, we collapsed the NT and +RNase datasets by bait prior to performing enrichment analysis. As expected of nucleic acid binding proteins, a number of the A3 PPI functional enrichments corresponded to RNA and ssDNA processes and complexes (**Figure 1E, Table S7**). While A3A shares a functional enrichment, *i.e.* protein folding, with A3B, A3D, and A3F, it is for different complexes and specific proteins (PDCL3, PHLP, TXND9, TCPA, TCPH, and TCPZ) related to protein folding functions (**Figure S4C**). The A3A-specific functional enrichment is for PDCL and TRiC/CCT in G-protein beta folding, and in a recent study it was shown that A3A interaction with the CCT complex inhibits its deaminase activity (58). Consistent with the proteomics results of the previous study, in our hands the CCT complex binding is specific to A3A and does not co-purify with other A3 proteins. A3C shares some functional overlap with other A3 proteins including mRNA and ncRNA metabolic processes (**Figure 1E**) (58). The top A3C-specific functional enrichment is the Replication Protein A (RPA) complex, a heterotrimeric complex that binds to ssDNA. Previously, it has been shown that RPA can interact with AID (59, 60) and recently it was shown that the RPA complex is co-opted by the L1 retrotransposon to facilitate integration and can also recruit A3 proteins to the site of integration, although A3C was not specifically studied (61). Interestingly, A3H-I and A3G, but not A3H-II interactors share a functional enrichment in tRNA processing (**Figure 1E, Figure S4C**). In addition, A3G-specific interactors are enriched for tRNA pseudouridine synthesis functions (**Figure 1E**). To our knowledge this is the first time A3G and A3H proteins have been linked to tRNAs and other tRNA editing enzymes. The A3B, A3D, and A3F proteins share a core group of interacting proteins and the main shared functional enrichment for this trio is in protein folding (**Figure 1E**). Unlike A3A, these three A3s do not have interactions with the CCT complex, their protein folding enrichment is related to interactions with prefoldin (PFD) complex subunits (**Figure 1E, Figure S4C**). Collectively, these results demonstrate that a number of A3 PPIs are mediated by RNA, and that while some PPIs are shared, a large number of interactors are A3-specific. These data demonstrate why it is important to systematically map interactions for all the A3 proteins under RNA-depleted and RNA-untreated conditions.

### The RNA -dependent and -independent A3 PPI Network

After identifying high confidence interactors for each bait, we wanted to identify which PPIs were condition specific, that is those that more strongly interact with specific A3 proteins in +RNAse, NT, or both conditions. A3 enzymes are known to interact with cellular RNA promiscuously, being identified as interacting with multiple RNA types in the absence of viral infection (62–64). Studies of A3 enzymes during HIV-1 infection and structural studies show that they prefer G-rich and A-rich sequences, but do not appear to have other specific requirements (65–67). Notably, recent cryo-EM structures have determined that A3G can form an interaction with the HIV-1 Vif protein that is dependent on RNA and protein contacts, thus providing a model for the RNA-dependent interactions identified in this study (67–69). Since RNA-dependent interactions were identified consistently across biological replicates, we hypothesize that there are both protein and RNA interactions that are taking place. Conversely, interactions that occur only in the +RNAse condition may occur due to RNA occluding the protein binding interface of the A3 or prey protein, now being exposed by the addition of RNAse A degrading the bound RNA. As with all AP-MS data, additional validation will be needed to ensure these interactions take place within proper cellular compartments and the context of living cells with intact signaling.

As expected, a number of the high confidence PPIs were RNA-dependent with 83 being identified only in the NT condition. In addition, we also identified 93 A3-interacting proteins only under +RNAse conditions. For a number of the PPIs classified as high confidence under one condition, they were pulled down and identified in both, sometimes just below our stringent high confidence threshold. Therefore, in order to more accurately discern conditional interactions from non-conditional interactions, we used a modified differential interaction score (DIS) similar to previous reports (31–33). The DIS maintains high stringency hit calling (BFDR<0.05; CompPASS wd 0.9 percentile per bait), while also recovering conserved interactions scoring below strict cutoffs for one condition but not for the other. Here, a DIS of 0 indicates that a prey was confidently purified under both NT and +RNAse conditions, whereas a DIS of +1 or −1 indicates that a prey interaction is specific to +RNAse or NT conditions respectively (**Table S8**). If we use a cutoff of |DIS| >0.5 indicating conditional change, we find that 163 PPIs are identified in both conditions, and that only 38 are preferentially identified in +RNAse conditions, and 35 are preferentially identified in NT conditions (**Table S8**). Combining NT and +RNAse PPI networks we capture 236 interactions between the eight A3 protein baits and 155 interacting proteins (**Figure 2**). It is important to note that since the SAINT BFDR accounts for more than just abundance, this is not an indicator of the relative abundance of an interaction, but a way to indicate if a PPI is an interactor in one or both conditions. For some PPIs identified as an interactor in both conditions, it may be that this interaction is stronger or more abundant in one condition. To this end, we performed MSstats analysis on each A3 bait PPI network, quantitatively comparing RNAse+ and NT conditions (**Figure S6, Table S8**).

**Figure 2.**
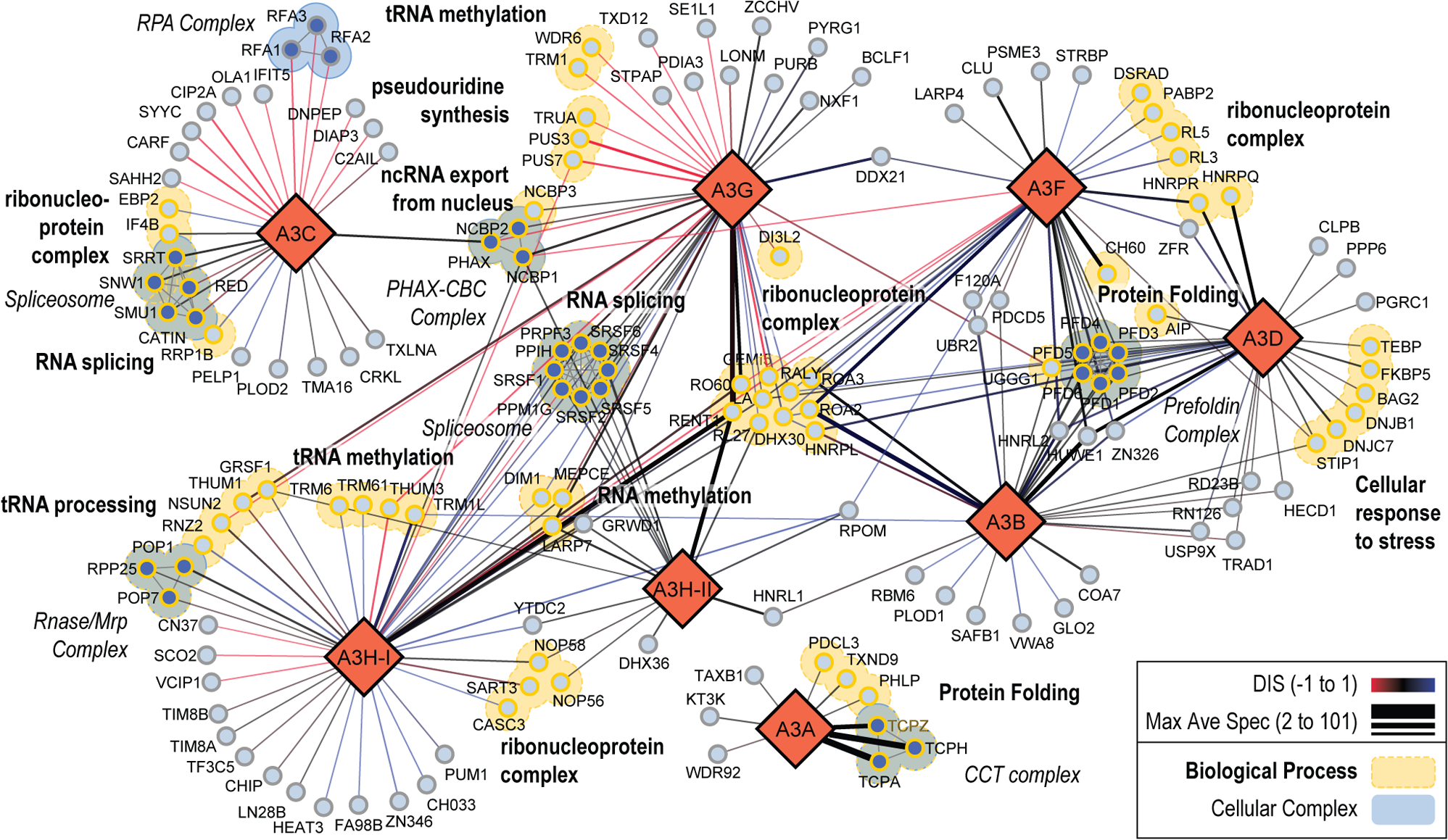
The A3 family RNA-dependent and RNA-independent PPI network. Protein interactions are depicted as edges drawn between two protein nodes with the edge thickness representing the maximum of the average spectral count (Max Ave Spec from 2 to 101) for a particular bait-prey pair. Edge color represents the DIS of the pairwise interaction from −1 to 1, with −1 being confidently identified only in NT (red), 1 being confidently identified only in RNAse+ conditions (blue), and 0 indicating similar interaction in both conditions (black). Gray edges are prey-prey interactions from known protein complexes. The A3 proteins are represented as large diamond nodes, and the identified prey proteins are represented as circular light blue nodes. Prey proteins involved in select enriched cellular complexes, are depicted as filled in blue nodes. Prey proteins involved in select biological processes are depicted with yellow colored borders.

In our differential A3 PPI network (**Figure 2**) edge width correlates to the average spectral count (AveSpec) and the edge color corresponds to the bait-prey DIS. For PPIs identified in both conditions, the edge width corresponds to the highest AveSpec (Max Ave Spec). Functional enrichments that corresponded to biological processes (BP) and cellular complexes (CC) were extracted and proteins in the full PPI network were mapped back to a single biological process or cellular complex. If a protein had more than one BP and one CC, the term with a larger population was selected to represent it in the network (**Table S8**). Several PPIs captured in this network have been described in previous studies, or have been reported in interaction databases (**Table S8**) (51, 52, 58, 70, 71), though we capture a number of novel interactors as well. The most interconnected A3 interacting proteins were RNA binding proteins including proteins involved in splicing, ribonucleoprotein complexes (RNP), and RNA modification. Specifically, components of the spliceosome interact with A3C, A3G, and both A3H haplotypes. Though A3C interacts with a different subset of spliceosomal components in an RNA dependent manner, A3G, A3H-I, and A3H-II interact with a core set of spliceosome factors in both conditions. To simplify, we can look at individual subnetworks for each A3 bait (**Figure S7**). The interconnectivity of a given prey is also mapped to each individual network (**Figure S7**). An in depth discussion of these results is presented as a Supplementary Discussion.

Although studies have identified the RNA-dependent interactions of A3G and A3F with RNP complexes previously [90,91], the RNA independent interactions and overlapping interactions with other A3 family members has not been examined in detail. Here we examine some of the RNA -dependent and -independent interactions that are identified in our PPI network. A3C, A3G, A3H-I and A3H-II capture and identify PPIs with functions in RNA splicing and spliceosome complexes. Though functionally similar, it is noted that the A3s showed distinct interactions with specific subunits. For example, A3C interacts with spliceosome proteins SRRT, SNW1, RED, SMU1, and CATIN, while A3G, A3H-I, and A3H-II do not. The DIS scores for SRRT, SNW1, and SMU1 were similar in NT and +RNAse, but RED and CATIN were stronger interactors in NT, indicating a potential dependence on RNA for the interaction. In contrast, A3G, A3H-I, and A3H-II share interactions with a separate spliceosome protein complex involving PRPF3, SRSF1, SRSF2, SRSF4, SRSF5, SRSF6, PPM1G, and PPIH, although not all three A3 proteins interact with all these components. Most interactions are shared between A3G and A3H-I, with only SRSF1, SRSF2, and SRSF6 being shared between A3G, A3H-I, and A3H-II. All of these interactions occurred under both conditions except SRSF1 and SRSF2 which were recovered for A3H-I only in the +RNAse condition. Additional PPIs were found with functions involved in tRNA processing, RNA methylation, and ncRNA export from the nucleus.

Similar to spliceosome components, PPIs with functions in tRNA processing and RNA methylation components were found to interact with multiple A3 proteins, primarily A3G and A3H-I, though many were non-overlapping between A3 datasets. A3G specifically interacted with tRNA processing and RNA methylation PPIs that were only observed in the NT condition (e.g., PUS7, PUS3, TRUA, TRM1, WDR6) whereas most PPI from this category with A3H-I were found in both NT or +RNAse, with less found with only NT (e.g., THUM3, PUS7) or +RNAse (e.g., RNZ2, TRM6, TRM61, DIM1). We validated the RNAse+ and NT condition specificity for two of these, PUS7 and RNZ2, in HEK293Ts overexpressing FLAG-tagged A3 proteins. Using co-immunoprecipitation (co-IP) immunoblots, we show that PUS7 is NT-specific while RNZ2 is RNAse+ specific, consistent with our MS results (**Figure S8A**). We also demonstrate the RNA-dependence of the PUS7 interaction by reciprocal co-IP of the endogenous PUS7 protein from HEK293T cells expressing A3G or GFP control (**Figure S8B**).

It was notable that A3B, A3D, and A3F all interacted with multiple members of the Prefoldin family of proteins (PFD1-6). The strongest interactions with A3B are PFD proteins, with PFD3, PFD5, and PFD6 being the most abundant, in both the NT and +RNAse condition (**Figure 2 and Figure S4**). PFD2 had the weakest interaction with A3B and A3F and PFD4 had an intermediate interaction strength with A3B, A3D, and A3F. In the case of A3D, there was no high confidence interaction with PFD2. The A3F interactions were 2-fold less strong than A3B and the A3D interactions were 3- to 4-fold less strong than A3B. However, all the interactions with the PFD proteins were usually similar for the NT and +RNAse conditions, except A3D and PFD4, which only occurred in the presence of RNAse A (**Figure 2 and Figure S4**). To validate the AP-MS results, we conducted reciprocal co-IPs where we used FLAG tagged PFD3 or PFD5 and immunoprecipitated 3×HA-tagged A3B, A3D, A3F, and A3G in the NT or +RNAse condition. A3G served as a negative control since it did not interact with any PFD family members in our AP-MS result, but is also a double domain A3 protein (**Figure 2 and Figure S4**). We observed that A3B and A3F had a strong interaction with PFD3 and PFD5 in the NT or +RNAse condition. Consistent with the AP-MS analysis, A3D showed less interaction, and A3G did not interact with PFD3 and PFD5 (**Figure 3, Figure S9**).

**Figure 3.**
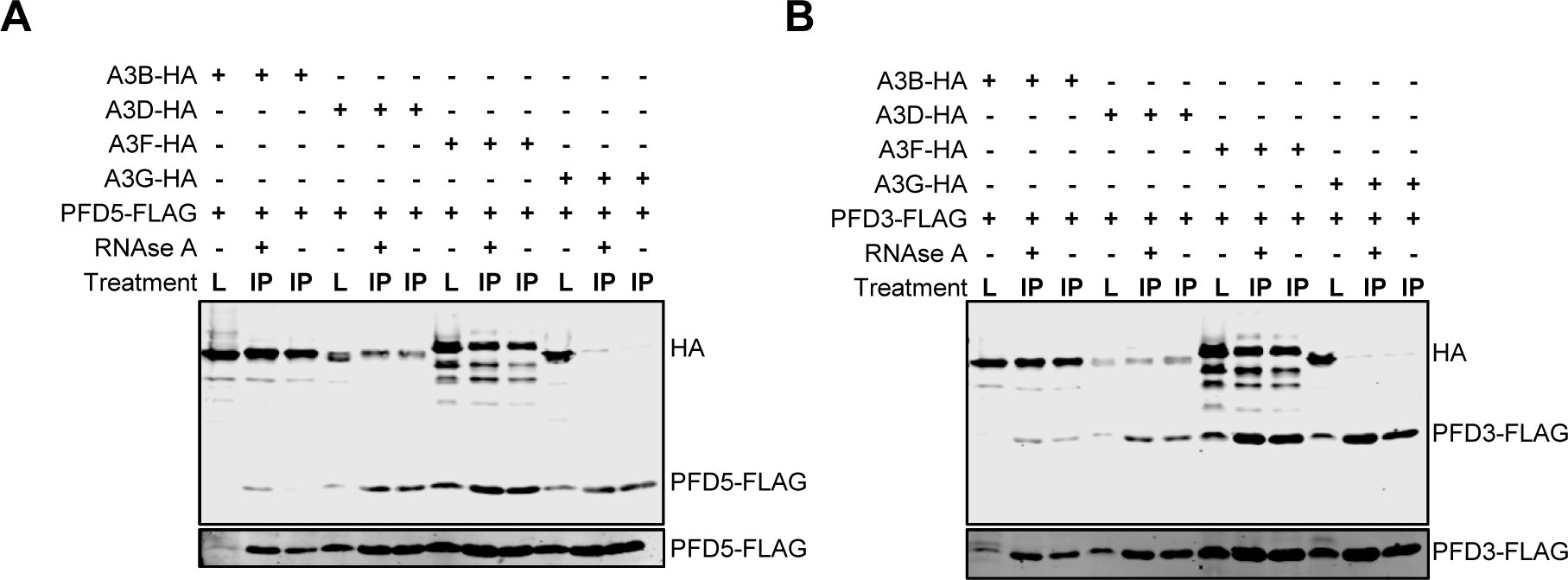
Multiple A3s interact with Prefoldin complex members. Co-IP of **(A)** PFD5 or **(B)** PFD3 with A3 enzymes. Cell lysates (L) from HEK293T cells were used for FLAG immunoprecipitation of PFD5- or PFD3-FLAG. Co-immunoprecipitating HA-tagged A3s were detected in either the NT or +RNAse immunoprecipitation (IP). Higher contrast PFD5 or PFD3 blots are shown below each blot to demonstrate the presence of PFD5 or PFD3 in each sample. Variable levels are due to PFD polyubiquitination and degradation (76). AP-MS results demonstrated an interaction of A3B, A3D, A3F, but not A3G with PFD3 and PFD5 with varying strengths. Consistent with this, the co-IP shows that in the NT or +RNAse condition, there are interactions between A3B, A3F, and A3D with PFD3 and PFD5, but not A3G. Purification control immunoblots are shown in **Figure S9.**

### A3B interaction with PFD5 inhibits PFD5-mediated degradation of cMyc

Prefoldins are present in all eukaryotes and Archaea and exist as a hetero-hexameric complex (72). The PFD subunits assemble into a β-barrel complex with six long tentacle-like coiled coils, that act as a molecular cochaperone to fold actin and tubulin monomers during cytoskeleton assembly, though each PFD also has distinct independent biological functions (72). The prefoldin complex acts on unfolded actin and α- and β-tubulin cotranslationally and delivers them to CCT posttranslationally (73). Despite this canonical role as a cochaperone, the Prefoldin subunits also shuttle between the cytoplasm and nucleus (PFD6, PFD2) or are predominantly nuclear (PFD5) and act on multiple DNA binding proteins (72). PFD3 localizes to the cytoplasm when solely expressed, but translocates to the nucleus in the presence of the Cullin 2 ubiquitin ligase substrate receptor, VHL (74). This PFD3-VHL interaction induces degradation of some DNA repair proteins such as the DNA mismatch repair protein MSH4 (75). All the PFD protein levels in cells are tightly controlled when not in a protein complex through polyubiquitination and degradation (76).

To study the functional implications of these interactions, we hypothesized that the role of A3s would not be to interact with the total PFD complex that has a role in protein folding but to interact with the individual PFD subunits, which have nuclear roles, often with DNA binding proteins. Since A3s have roles in modifying DNA, we investigated if any PFDs and A3s would have any overlap (10, 77, 78). In particular, A3s also have deamination-independent roles that solely rely on their ability to bind nucleic acids, such as restriction of retrotransposons, inhibition of viral polymerases, and A3A was recently identified to induce genomic instability in pancreatic cancer by a deamination independent mechanism (4, 5, 13). Thus, we also considered that the protein interaction network could reveal additional deamination independent functions for the A3 family.

PFD5 (also called MM-1) is a well characterized tumor suppressor since it acts as a co-repressor of E-box dependent transactivation activity of cMyc (79). PFD5 can bind the N-terminal region of cMyc and repress transcriptional activity in three different ways. This can be through PFD5 recruiting the HDAC1-mSin3 complex to inhibit chromatin remodeling, monoubiquitinated PFD5 recruiting a Skp2-ElonginC-ElonginB-Cullin2 complex to induce cMyc degradation, or PFD5 and the Egr-1 repressor binding and downregulating the wnt4 gene, that would otherwise target cMyc gene expression (72). The most direct way that PFD5 could inhibit cMyc is through the recruitment of the Skp2-ElonginC-ElonginB-Cullin2 complex for ubiquitination and proteasomal degradation of cMyc (**Figure 4A**) (80). A3B bound to PFD5 could potentially block the interaction with cMyc and disrupt PFD5 tumor suppressor activity (**Figure 4A**). This would be consistent with recent results that show A3B mRNA, but not deamination activity is associated with cancer (81), suggesting that A3B has a deamination independent effect on cell proliferation (81). Moreover, this would be consistent with cMyc usually being dysregulated but not lost in many cancers (82). To test whether A3B interaction with PFD5 inhibited its interaction with cMyc, we co-purified PFD5-FLAG, A3B-HA, and cMyc in the presence of the proteasome inhibitor MG132 (**Figure 4B-C**). The PFD5-FLAG was co-expressed in HEK293T cells with A3B-HA and cMyc alone or together. The FLAG tag was immunoprecipitated and the resolved proteins were blotted for cMyc, HA, and FLAG. Both cMyc and A3B-HA interacted with PFD5-FLAG independent of each other. In addition, using 1 µg of each plasmid, we observed that PFD5 could interact with both cMyc and A3B-HA at the same time (**Figure 4B-C**). However, there was less cMyc in the co-IP when 1 µg of cMyc compared to 4 µg of cMyc was co-transfected with 1 µg A3B-HA (**Figure 4B-C**). When 4 µg of cMyc was co-transfected with 2 µg A3B-HA, the level of cMyc in the co-IP decreased further than when co-transfected with 1 µg A3B-HA (**Figure 4B-C**). Altogether these data suggest a competitive interaction of cMyc and A3B with PFD5-FLAG. To determine if this interaction could disrupt PFD5-mediated degradation of cMyc we coexpressed PFD5-FLAG, A3B-HA, and cMyc in the absence and presence of MG132. We observed that in the presence of PFD5-FLAG, the cMyc is decreased by 1.5-fold (**Figure 4D-E**). When A3B-HA is coexpressed with PFD5-FLAG and cMyc, the cMyc is 3.3-fold greater than cMyc alone (**Figure 4D-E**). This large increase in the cMyc by A3B-HA may be due to the added protection from endogenous PFD5 in the HEK293T cells. Consistent with this is that in the presence of MG132, the cMyc levels in cells is equivalent to that in the presence of A3B-HA. Further, the PFD5-FLAG mediated degradation of cMyc is blocked in the presence of MG132 and the recovery of cMyc protein levels in the presence of MG132 and A3B-HA is 7.5-fold above cMyc expression alone. The higher rescue of cMyc in the presence of MG132, compared to its absence suggests that A3B alone cannot fully compete for PFD5 binding with cMyc, consistent with the co-IP data (**Figure 4D-E**).

**Figure 4.**
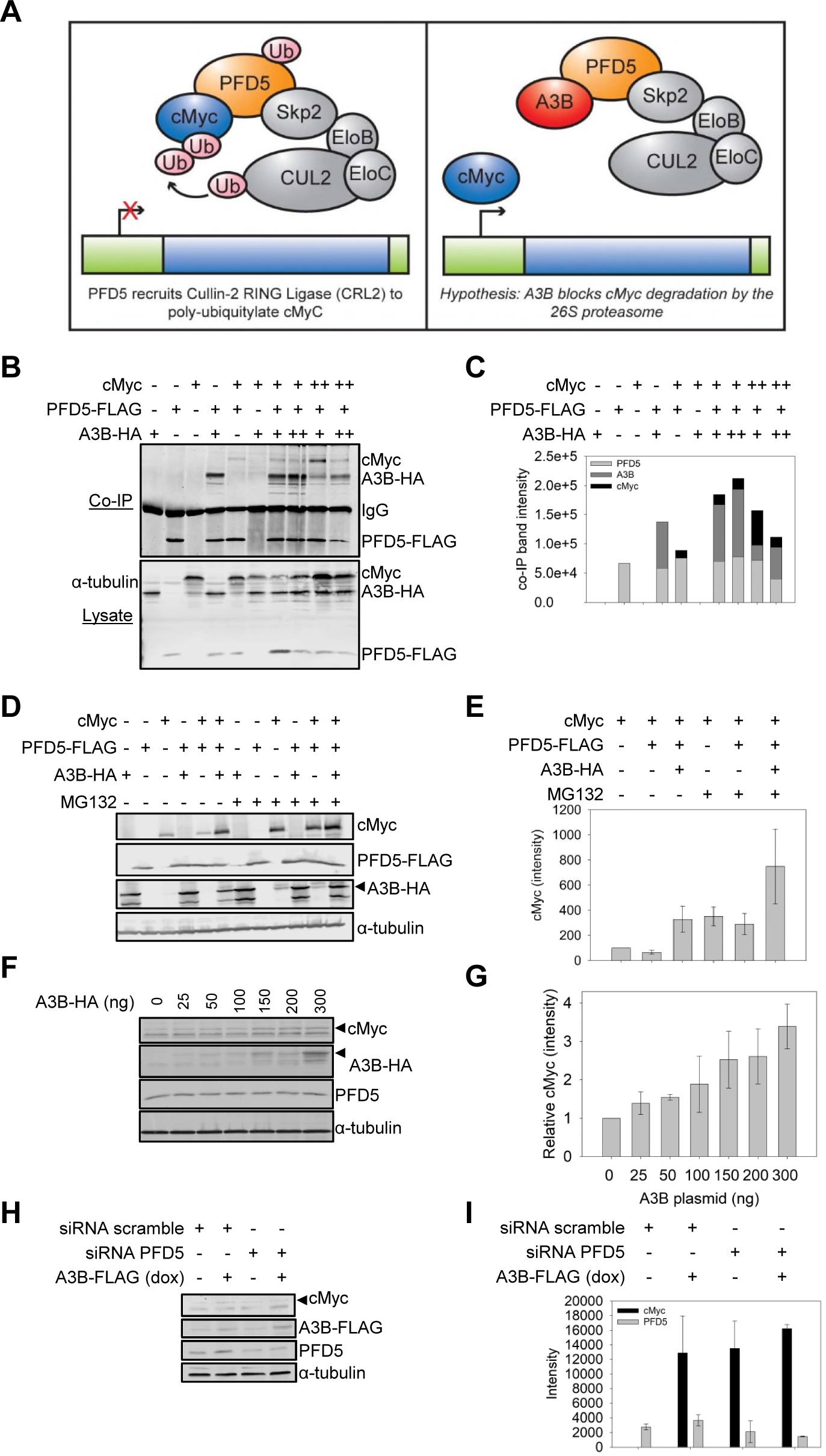
A3B inhibits PFD5-mediated degradation of cMyc. **(A)** Left: Diagram of tumor suppressor PFD5 recruiting Cullin-2 RING ubiquitin ligase for cMyc ubiquitylation and degradation. Right: A3B interacts with PFD5 leading to the hypothesis that A3B can disrupt this degradative pathway and lead to A3B-mediated, but deamination independent, dysregulation of the cell cycle. **(B-C)** Co-IP experiments from HEK293T cells treated with 12.5 μM MG132 for 16 hr demonstrated that A3B-HA and cMyc each interact with PFD5-FLAG. Using combinations of transfections of low A3B (1μg, +), high A3B (2μg, ++), low cMyc (1μg, +), or high cMyc (4μg, ++) demonstrated that the presence of A3B resulted in less co-purification of cMyc with PFD5 in a concentration dependent manner. **(C)** is the quantification of **(B)**. **(D-E)** This interaction was functional and resulted in more cMyc in cells in the presence of A3B-HA, when PFD5-FLAG was also present. **(E)** is the quantification of three independent blots, with a representative blot shown in **(D)**. **(F-G)** A3B-HA regulation of endogenous PFD5 occurs in the MCF7 breast cancer cell line. MCF7 cells were transfected with increasing amounts of A3B-HA. The endogenous cMyc detected increased with the amount of A3B-HA protein in cells. The cMyc was quantified relative to the amount of α-tubulin in each lane and normalized to the no A3B condition (value of 1). Quantification is plotted in **(G)** with error bars that represent the standard deviation from two independent experiments. A representative blot shown in **(F)**. **(H-I)** MCF7 cells with doxycycline inducible A3B-FLAG were transfected with siRNA scramble or siRNA PFD5 to knock down endogenous PFD5 in the absence or presence of doxycycline. **(H)** The effect on the endogenous cMyc was determined by immunoblotting. **(I)** The intensity of the cMyc and PFD5 was quantified and is shown with the error bars that represent the standard deviation from two independent experiments.

To determine if this relationship occurs in a cell type relevant to cancer, we used the MCF7 tumorigenic breast cancer cell line that is estrogen receptor positive and does not express A3B in the absence of estrogen. cMyc amplifications are found in 44% of ER positive breast cancers and cMyc can also affect cancers by resisting degradative pathways to maintain protein stability in cells (82). In this experiment we transfected increasing amounts of A3B-HA expression plasmid in the absence of MG132 and detected the effect on endogenous cMyc and PFD5 (**Figure 4F-G**). We quantified the amount of cMyc relative to the level of α-tubulin to normalize any differences in total protein.The steady state cMyc protein levels increased to a maximum of 2.5 to 3.4-fold when 150 to 300 ng of A3B plasmid was transfected (**Figure 4F-G**). The PFD5 levels remained constant. To confirm that this relationship was dependent on the interaction between A3B and PFD5, we conducted experiments in an MCF7 cell line that expresses doxycycline (dox) inducible A3B-FLAG and transfected in siRNA against PFD5 or a scrambled siRNA (**Figure 4H-I**) (40). The siRNA resulted in a 2-fold decrease in PFD5 **(Figure 4H-I**). The data confirmed that in the presence of PFD5 (siRNA scramble), cMyc protein levels are low unless A3B is present (siRNA scramble, +dox) (**Figure 4H-I**). However, when PFD5 expression was knocked down (siRNA PFD5), cMyc protein levels were high regardless of the A3B protein levels (dox- and dox+) (**Figure 4H-I**).

## DISCUSSION

Despite A3 enzymes being highly studied for their response to viral infection and role in cancer, there have been very few studies on interacting proteins and in some studies it was unclear whether interactions were mediated by RNA or not. Here we determined the protein-protein and protein-RNA mediated interactions for eight A3 enzymes. The A3 family has highly similar amino acid sequences due to being formed through duplication events, however, several distinct interactions suggest that A3 activity is not fully redundant and may have deamination-independent functions.

Shortly after the discovery of the A3 family, the family was divided into three Z-domains. This resulted in Z1, Z2, and Z3 groups, of which A3H is the only Z3 member. Some A3s with two deaminase domains have a combination of Z1 and Z2. We found that there were more interactions shared between similar Z-domains, than less related Z-domains, but there were still distinct interactions for all A3s. Even A3H-I and A3H-II which are identical except for three amino acids (105, 121, and 178) had distinct interactions. Perhaps most surprising was that A3G that had amino acid similarities with A3B, A3D, and A3F shared very few PPIs with these 3 A3s that had multiple shared PPIs. Rather A3G (Z2-Z1 domains) clustered with the Z3 group (A3H) under NT and by itself under +RNAse conditions respectively. Even A3A that has a high amino acid similarity to the A3B CTD shared no interactions with A3B or any other A3.

Prior to this study, protein network analyses found similar interactions for A3G and A3F with RNA binding proteins, which overlap with our protein interaction network (51, 52, 62). However, an important difference with our study is that the previous interactions were primarily detected with the NT condition and the authors concluded that there are few significant interactions in the +RNAse condition. In contrast, here we find that many of the RNA binding protein interactions of A3G and A3F occur in both the NT or +RNAse conditions and that A3F interacts with other types of proteins, such as those involved in protein folding. Although Kozak et al. determined that many of these RNA binding proteins are not incorporated into HIV-1 virions, the only function identified for A3G and A3F at the time was restriction of HIV replication and no other functions based on the interacting proteins were proposed (51). The interactions previously published may be fewer or different due to those studies conducted in T cell lymphoblastic cell lines. However, it is now known that A3s can be upregulated in any epithelial cell after a virus infection and during cancer their mRNA expression has been found in multiple tissues (83–86). Thus, the potential interactions of A3 enzymes is more diverse than previously thought.

The protein interaction network of A3C revealed some potential functions. Although A3C is highly expressed in T cells (42) it has minimal restriction activity against HIV-1 (87, 88), but can restrict the retroelement LINE-1 (87, 89). However, the interaction of A3C with CATIN/CACTIN suggests a possible reason for the high expression of A3C. CATIN/CACTIN is a negative regulator of Toll-like receptor (TLR) activity and interaction with A3C may de-repress TLR activity ensuring a proper immune response (90). Additionally, the interaction of A3C with the complex RED/IK and SMU1 that is required for mRNA splicing, in particular splicing of influenza A virus NS1 pre-mRNA, may act to suppress splicing and exert an antiviral response (91). Since A3C catalytic activity is low, it is fitting that these potential functions do not rely on deamination activity (92). Although there is a more active version of A3C in the human population, A3C S188I, this form is only found in 10% of people of African descent, suggesting that the primary role of the common A3C is deamination independent (87).

The interaction of A3G and A3H with tRNA binding and modification proteins suggests a role in tRNA biology. Although deamination of adenosine to inosine in tRNAs is a well characterized deamination event, cytosine deamination has not been documented (93). Based on the ability of A3 enzymes to regulate HIV-1 reverse transcriptase by binding to the RNA template or the enzyme itself, the roles of A3G and A3H may be to regulate activity of other enzymes by binding the tRNA (94, 95). This may relate to an antiviral role if A3G and A3H can temporarily slow or shut down protein synthesis during a viral infection, which would facilitate immune clearance.

Overall, the interactions with RNA binding proteins by all A3s may also be a mechanism to inhibit LINE-1 retrotransposition. Although most A3 enzymes inhibit LINE-1 movement, only A3A has been shown to do this by a deamination-dependent mechanism (96, 97). Other A3s have a deamination-independent mode of restriction that has been poorly characterized. A3C and A3D have been reported to interact with the LINE-1 protein ORF1p as a mechanism of inhibition, but it is not known how A3B, A3F, A3G, and A3H can restrict retrotransposons (89, 98, 99). The protein interaction network suggests many possibilities of disrupting LINE-1 mRNA transport. For example, A3G interacts with NXF1, which facilitates mRNA export from the nucleus and is also used by LINE-1 (100, 101). Further confidence in this type of role comes from interaction of A3G, A3H-I, and A3H-II with SRSF complex proteins (SRSF1, 2, 4, 5, and 6) that function as export adapters for the TAP/NXF1 nuclear export pathway (102). A3F, A3G, and A3H-I also interact with RO60 that binds to endogenous retroelements, such as Alu (103). ROA3 that plays a role in cytoplasmic trafficking of RNA interacts with A3F and A3B (104). These interactions suggest that A3 enzymes may be able to affect the mRNA transport of LINE-1 or Alu, disrupting their ability for nuclear export or import of their mRNA.

One of the key interactions we identified was with the prefoldin family of proteins and A3D, A3F, and A3B. Although these proteins are well-known chaperone proteins, they also have individual roles. However, it is also interesting that A3A interacts with proteins of another chaperone complex, the CCT complex that includes TCPA, TCPZ, and TCPH. For A3A, it was found that interaction with the CCT complex inhibits A3A deamination activity, which may be a mechanism to protect from unwanted deamination of genomic DNA (58). We did not find that the PFD proteins specifically inhibited the activity of A3D, A3F, or A3B (data not shown). Rather, we found that A3B could inhibit the normal tumor suppressor role of PFD5 to induce degradation of the oncogene cMyc. Further studies are needed to determine the long term impact of A3B-induced increases of cMyc in cells. Although the discovery of the CCT complex inhibiting A3A activity was initially thought to protect the genomic DNA from damage, it was found that tumors with an A3A mutation signature also harbor CCT complex mutations that may derepress A3A activity (58). Nonetheless, A3A has also been found to have a deamination independent role in causing genomic instability in pancreatic cancer, although the mechanism is not known (13). For A3B, we observed that competitive binding for PFD5 with cMyc disrupts the cMyc degradation pathway, which can promote cellular proliferation.These data demonstrate that despite there being clear mutation signatures of A3 enzymes in multiple cancers, they may be acting in a deamination-independent manner, which should be considered alongside development of catalytic inhibitors for cancer treatments (105).

The data presented here suggest new potential roles of A3 enzymes. Since the discovery of A3 enzymes it was known that most of them exist in RNPs. However, the identity of the interacting proteins in these complexes was never determined for all A3 enzymes, which led to difficulties in ascribing a function to these RNPs. These data open several new avenues of investigation into their roles in tRNA maturation, RNA splicing, and ncRNA. In addition, we have for the first time identified high confidence protein-protein interactors of A3 enzymes suggesting additional functionality by which they contribute to known functions of viral and retrotransposon restriction and oncogenesis.

## ACKNOWLEDGEMENTS

We thank Tomas Pelletier for assistance on this project. We also thank additional members of the Krogan and Chelico groups for helpful comments and discussion. This research was supported by NIH/NIAID training grant F32AI127291 (to R.M.K); NIH/NIAID funding for the HIV Accessory & Regulatory Complexes (HARC) Center (P50 AI150476, N.J.K.); Saskatchewan Health Research Foundation Postdoctoral Fellowship Funding (A.G. and M.G.); College of Medicine Research Award (A.K.A.S.); College of Medicine Graduate Student Award (A.S.); Natural Science and Engineering Council of Canada (RGPIN-2016-04113, L.C.), and Canadian Institutes of Health Research (PJT-159560, L.C.).

## DATA AVAILABILITY

All data are contained within the manuscript and are available via ProteomeXchange.

This article contains supplemental data.

