## Supplementary figures and images for "Protein interaction map of APOBEC3 enzyme family reveals deamination-independent role in cellular function"

### Supplementary Figures

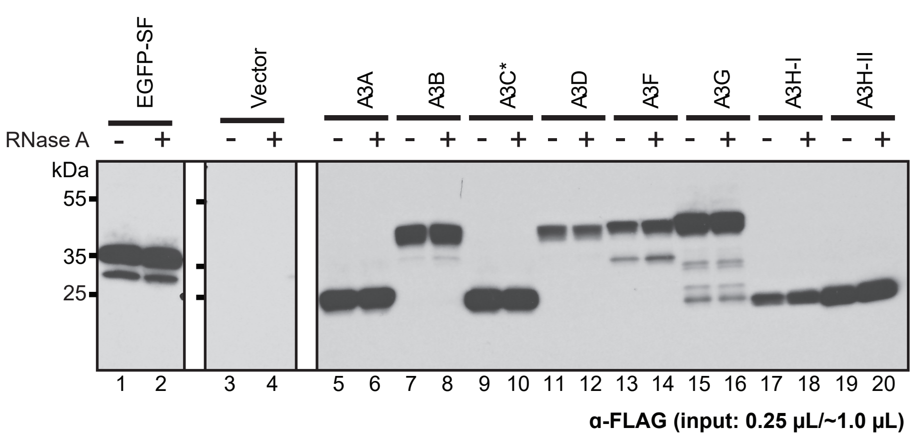

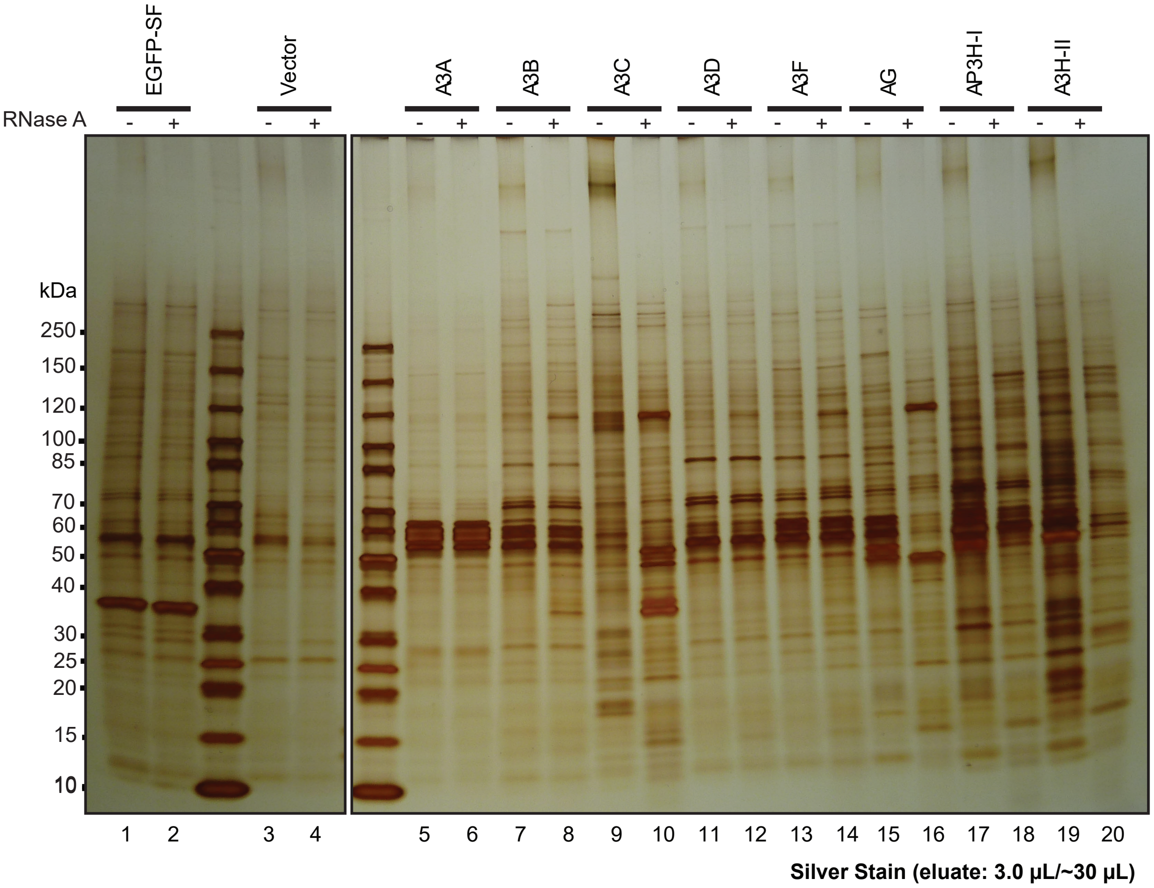

#### SAINT BFDR < 0.05 Thresholded PPI Network

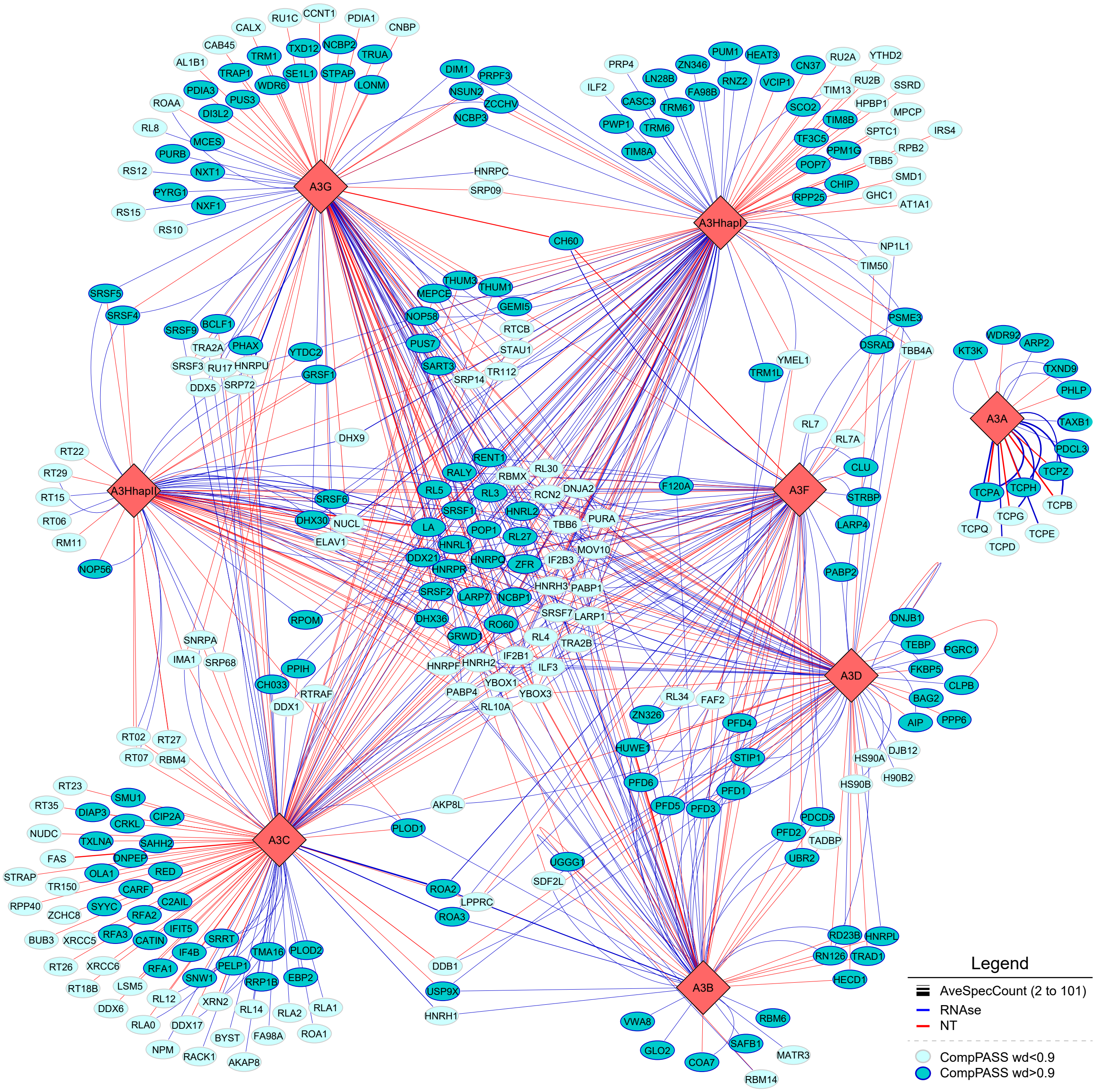

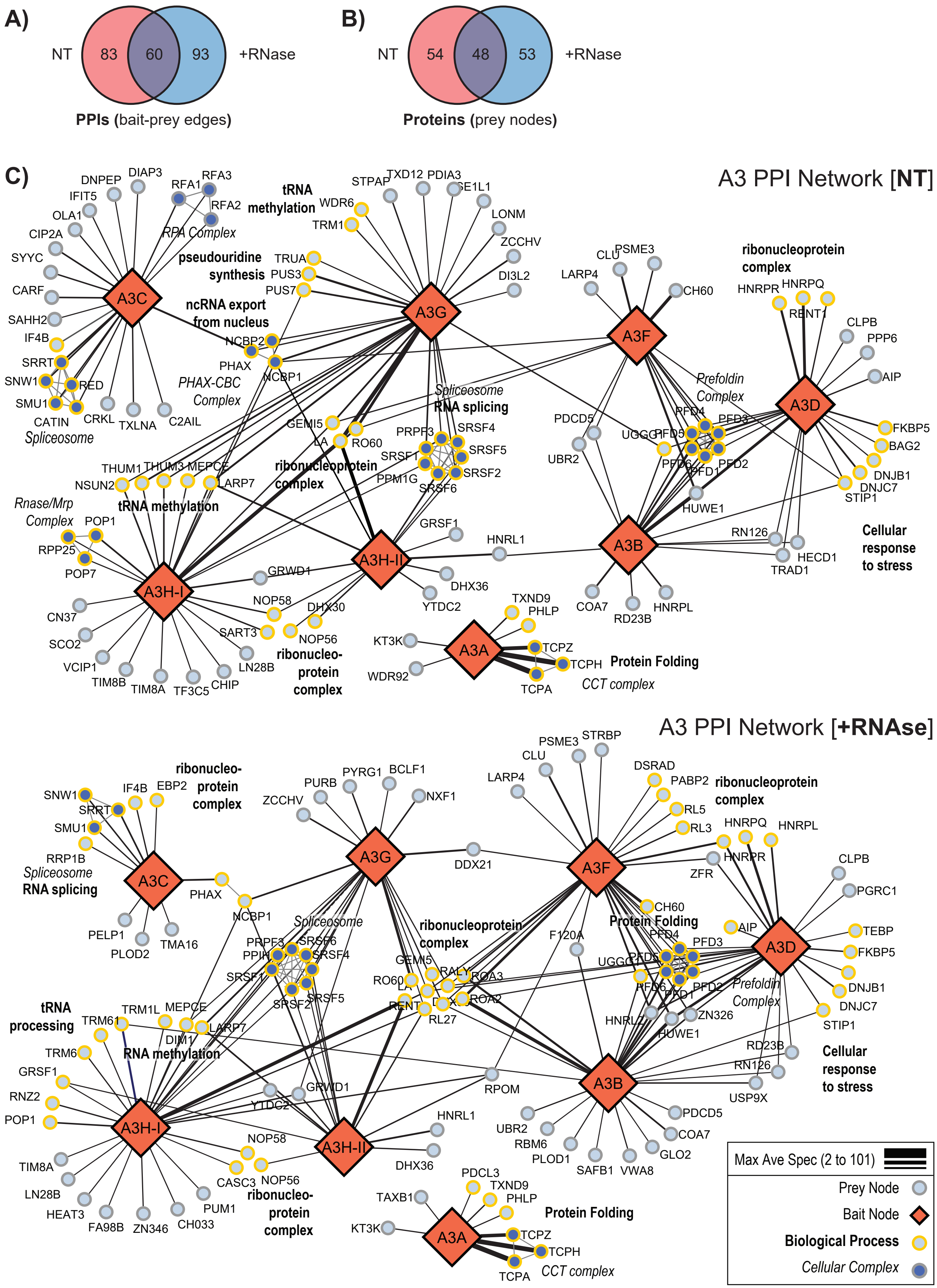

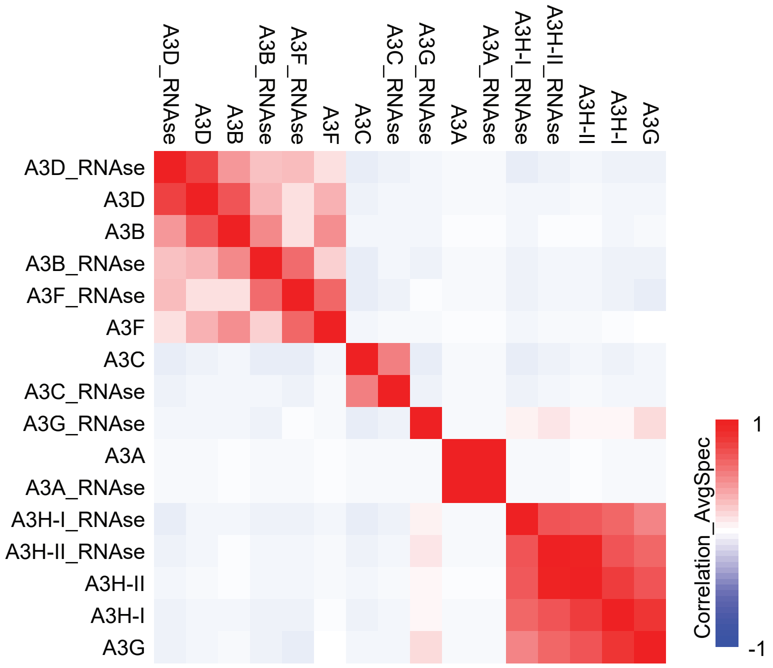

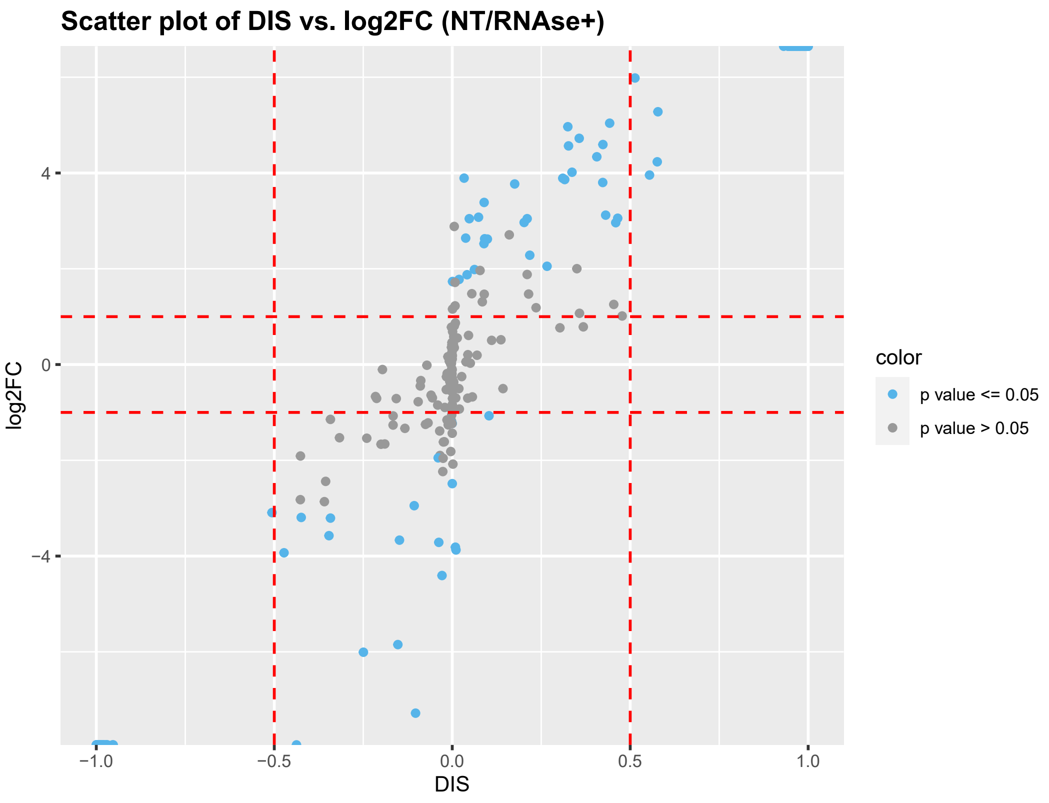

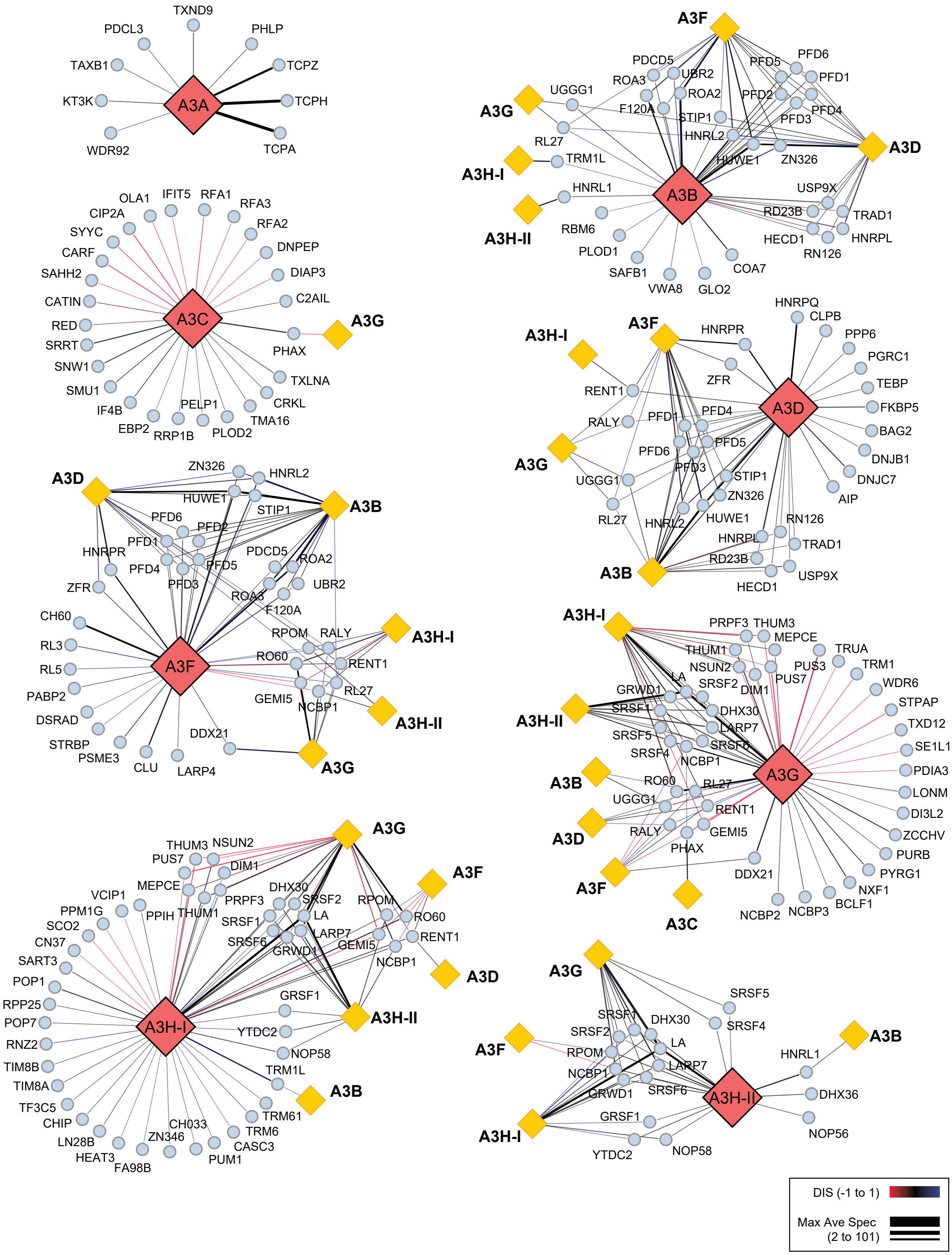

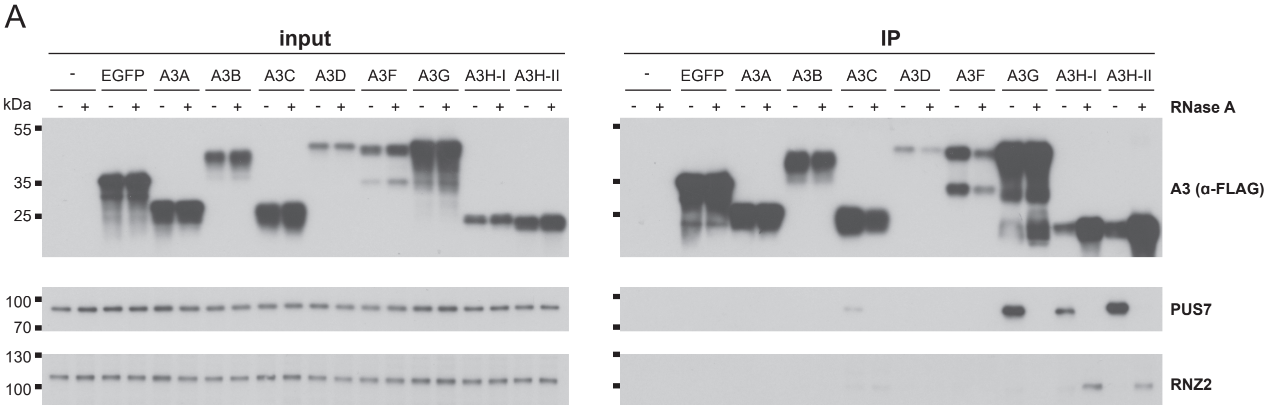

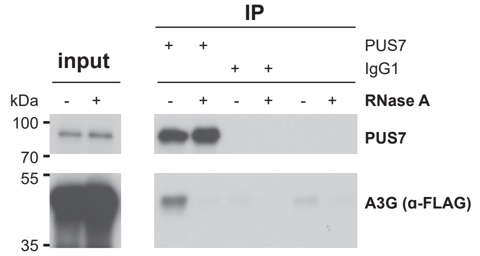

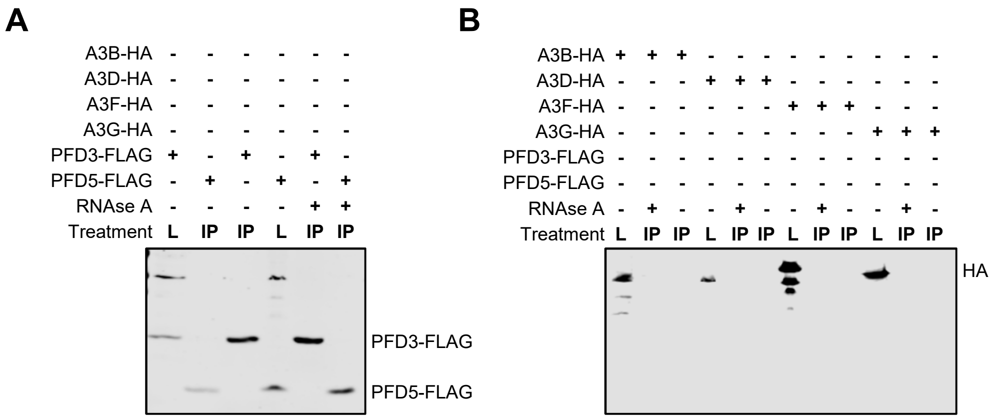
